# Predator-Prey Interactions Give Rise to Scale-Invariant Ecosystems

**DOI:** 10.1101/2022.07.03.498605

**Authors:** Mohammad Salahshour

**Affiliations:** Max Planck Institute for Mathematics in the Sciences, Leipzig, Germany; Max Planck Institute of Animal Behavior, Radolfzell, Germany; Centre for the Advanced Study of Collective Behaviour, University of Konstanz, Konstanz, Germany; Department of Biology, University of Konstanz, Konstanz, Germany

## Abstract

A large body of empirical evidence has shown that scaling laws and power-law distributions are ubiquitous features of ecological systems. However, it is not clear what factors give rise to such universal regularities and how scaling laws and scale-invariant distributions relate to each other. Here, I show scaling laws can be a simple consequence of scale-invariant distributions, and both result from simple commonalities of apparently different ecosystems. I introduce a simple model of predator-prey interactions in which predators and prey move on a two-dimensional space in search of resources that they use to survive and reproduce. The model predicts predator-prey systems can be found in different phases: as primary resources increase, the system exhibits a series of consecutive transitions to different phases with equilibrium dynamics and top-down control of the food web, non-equilibrium dynamics with bottom-up control, or unstable dynamics. While unstable predator-prey dynamics can result in a homogeneous environment by enrichment of resources, resource heterogeneity restores the stability of the system. Scale-invariant group size distribution and a rich set of scaling laws result in the non-equilibrium phase. By developing a general theory, I argue scaling laws can result from the scale-invariance of group size distributions under broad conditions. While some of the scaling laws predicted by the model await empirical examination, consistently with a recently discovered empirical pattern, the model shows predator abundance and prey production show a sublinear scaling with prey abundance. The model links the nature of the control in the food web, prey’s and predators’ behavioral responses to each other, and their life histories. While in small densities, mass-action law holds, and a weak density dependence and top-down control follows, in large densities, a shelter effect - the benefit of living in groups for preys - plays a role and makes prey production non-invariant to density fluctuations. Due to higher density fluctuations in higher densities, prey per-capita production in large densities decreases. This leads to a longer lifespan and lower population turnover of prey, and a scale-invariant ecosystem with bottom-up control in large densities.

---

Scaling laws and power laws are ubiquitous features of many biological and ecological systems [1–3]. These laws arise across different levels, from the individual level, such as metabolic scaling laws pertaining to individuals’ attributes [3, 4], to the population level, such as Taylor’s power law relating populations’ fluctuation and abundance [5, 6], or even in ecosystems and evolutionary level [7–10]. Among the scaling laws at work in ecosystem level, recently, a curious scaling law, according to which both predator abundance and prey production show a sub-linear power-law dependence on prey abundance, with an exponent close to 3*/*4 is argued to be at work across different ecosystems from terrestrial to marine communities [11]. Given the stark differences in composition and underlying interactions in these ecosystems, the emergence of such universal patterns may suggest some underlying scale-invariant properties are at work in ecosystems that originate from simple commonalities shared by diverse ecosystems. Indeed signatures of scale-invariance, such as scale-invariant group size distributions, have been observed in many biological populations [12–16]. Yet, it is not clear how such distributions relate to scaling laws and what factors give rise to these scale-invariant properties? To address this question, here I introduce a simple agent-based model of predator-prey interaction in which preys and predators move on a two-dimensional space in search of resources, which they use to pay for their metabolic costs and reproduce. Preys use primary resources regenerated in the system at a constant rate. Predators, on the other hand, reach resources by catching prey.

As the primary resources increase, the model shows different phases in which predators do not survive, prey and predators coexist in equilibrium and the system exhibits top-down control of the food web, and a non-equilibrium state in which prey and predators exhibit highly aggregated dynamics in the form of traveling waves and the food-web shows bottom-up control. These observations shed theoretical light on empirical arguments for the existence of such different phases in predator-prey systems [17–19], and the conditions under which they may show top-down or bottom-up control [20–22]. The predator-prey dynamics become unstable in too rich environments, resulting from the too fast exponential growth of predators acting faster than the regulatory feedback from prey to predator abundance. The instability of the predator-prey dynamics in resource-rich environments, coined the paradox of enrichment, has been argued theoretically [23, 24], and observed experimentally [17, 18]. As the model shows, in contrast to what has been suggested before [25], although spatial structure increases the stability, it fails to restore the stability of the system fully. However, as I show here, resource heterogeneity - a nonuniform distribution of resources in space - restores the stability of the system.

A rich set of scaling laws emerges in large densities, where the system exhibits non-equilibrium behavior and bottom-up control. Several variables, including prey and predator densities, density fluctuation, birth and death rates, and the maximum group size of prey and predators, obey a power-law relation with the resource regeneration rate. This implies these variables obey a power-law relation with respect to each other. This finding reproduces some of the scaling laws known to empirically hold in many ecosystems and suggests that many more such universal patterns can be at work awaiting empirical examination. For instance, the model predicts predator and prey density fluctuations scale as a function of each other with an exponent close to 1 in large densities, which recovers Taylor’s law with highly aggregated dynamics [5]. In addition, predator density and prey production obey a scaling relation as a function of prey density with a sublinear exponent. Furthermore, the model predicts group size distribution obeys a power law with a sharp cut-off. The cut-off, the largest group size, shows a scaling law with respect to resource regeneration rate. I will develop a general theory extending beyond predator-prey interactions to argue that these two conditions can give rise to a scale-invariant ecosystem exhibiting power-law dependence of population densities and highly aggregated dynamics. Given that the scale invariance of group size is argued to hold in many biological populations [12–16], the theory developed here suggests similar scaling laws may be at work in other contexts where populations coexist and interact.

The model also sheds light on one of the most fundamental questions in ecology, the direction of control in food webs [20–22], by establishing a link between the nature of control, prey and predator life histories, their collective dynamics, and behavioral responses to each other. In large densities with bottom-up control, prey’s lifespan increases, and their per-capita reproduction decreases with density, while predators’ lifespan decreases and their per-capita reproduction slightly increases with density. As the birth rate of predators and preys scale almost linearly, a higher lifespan of prey in rich and dense environments implies a sublinear scaling of predator abundance with respect to prey abundance. On the other hand, in small densities with top-down control, predators’ lifespan shows little density-dependence, and their per-capita reproduction increases with density. Preys’ lifespan decreases, and their per-capita reproduction increases with density. These observations link the top-down or bottom-up nature of control in food webs with the life histories of prey and predators. I will argue that a sublinear scaling in large densities and the nature of the control in the food web results from behavioral responses of prey and predators to each other. In large densities, a shelter effect - the benefit of living in large groups which can result from preys’ or predators’ behavioral responses to each other - plays a role, and the system exhibits highly aggregated non-equilibrium dynamics. In contrast to a simple mass-action law underlying Lotka-Volterra models [26–29], the shelter effect makes the resource flow to predators non-invariant to, and a decreasing function of density fluctuations. Due to higher density fluctuations in higher densities, resulting from Taylor’s law with highly aggregated dynamics, prey per capita production in large densities decreases. This leads to a longer lifespan and lower population turnover of prey in large densities and a sublinear scaling of predator-prey densities. On the other hand, in small densities, the shelter effect does not play a role, mass-action law holds, preys exhibit weak density dependence, and the system shows top-down control. These findings point toward the importance of preys’ and predators’ behavioral responses to each other in determining the nature of the control and dynamics of predator-prey interactions.

In the model, preys and predators move on a grid of linear size *L*. Each agent *α*, prey or predator, has an internal resource, *1*_*α*_, which increases by gaining resources, *κ*_*α*_, and decreases by a constant rate, *η*, due to paying for metabolic costs. Thus, the internal resource of agent *α* at time *t* evolves according to:

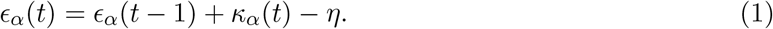

Preys acquire resources by consuming primary resources, which are regenerated on each site ***r***, with a constant rate *λ*(***r***). Basal resources available on a site are divided among all the prey visiting a site. Predators capture preys to gain resources. When a predator visits a site where prey lives, the predator captures a prey with probability *q*. When capturing prey, the internal resource of the predator increases by an amount *γ*. Preys and predators die if their internal resource reaches zero and reproduce if their internal resource reaches a threshold *d*. When reproducing, the individual produces an offspring, and the resources of the individual is divided between the individual and its offspring. If it happens that one of the species goes to extinction, assuming there is migration, one individual of that species is added to a random location on the lattice.

In the following, I will consider both a homogeneous environment, in which the resource on all the sites are regenerated with a constant rate *λ*, and a heterogeneous environment, in which the resource regeneration rate depends on the site. For this purpose, in the main text, I will consider a linear distribution of resource regeneration rate in which the resource regeneration rate *λ*(***r***) on a site ***r*** = (*x, y*) is given by *λ*(***r***) = 2*λx/L*. The case of binary resource distribution is considered in the Supplementary Information, and shows a similar phenomenology (see the Section Methods and Supplementary Notes. 6). For the spatial structure, both a nearest neighbor square lattice with linear size *L*, fixed boundaries, and von Neumann connectivity and a fully connected network, where individuals can jump to any site in the system, are considered.

The phase diagram of the model in the *λ − γ* plane and the densities of preys and predators as a function of *λ* for *γ* = 0.5 are plotted in, respectively, Fig. 1(a) and Fig. 1(b). With too low resource regeneration rates, predators do not survive. In this regime, the stationary density of prey is determined by equating resource regeneration and resource consumption, *λ* = *ηρ*_*p*_, which gives *ρ*_*p*_ = *λ/η*. By increasing *λ*, a phase transition above which predators survive occurs (blue circle). The transition occurs when the flow of resources to predators is larger than the resource consumption rate. A naive mean-field argument predicts the transition occurs at *λ*^*∗*^ = *η*^2^*/γq* (by equating resource flow to predators, *γqρ*_*p*_, and predator resource consumption *η*, and using *ρ*_*p*_ = *λ/η*, see Section Methods), in good agreement with simulation results (Fig. 1(a)). Above the transition line, the system reaches an equilibrium state where preys and predators coexist and no collective movement emerges (See the Supplementary Videos and Supplementary Note. 1 and 2 for the dynamics of the model in this regime). In this region, the density of prey shows little variation and slightly decreases with increasing *λ*. Instead, higher resources available in the system are transferred to predators, and their density increases by increasing *λ*. A simple mean-field argument predicts the density of preys in the equilibrium to be equal to *η/*(*γq*), with fair agreement with simulation results (see Fig. 1(c) and Section Methods).

**Figure 1:**
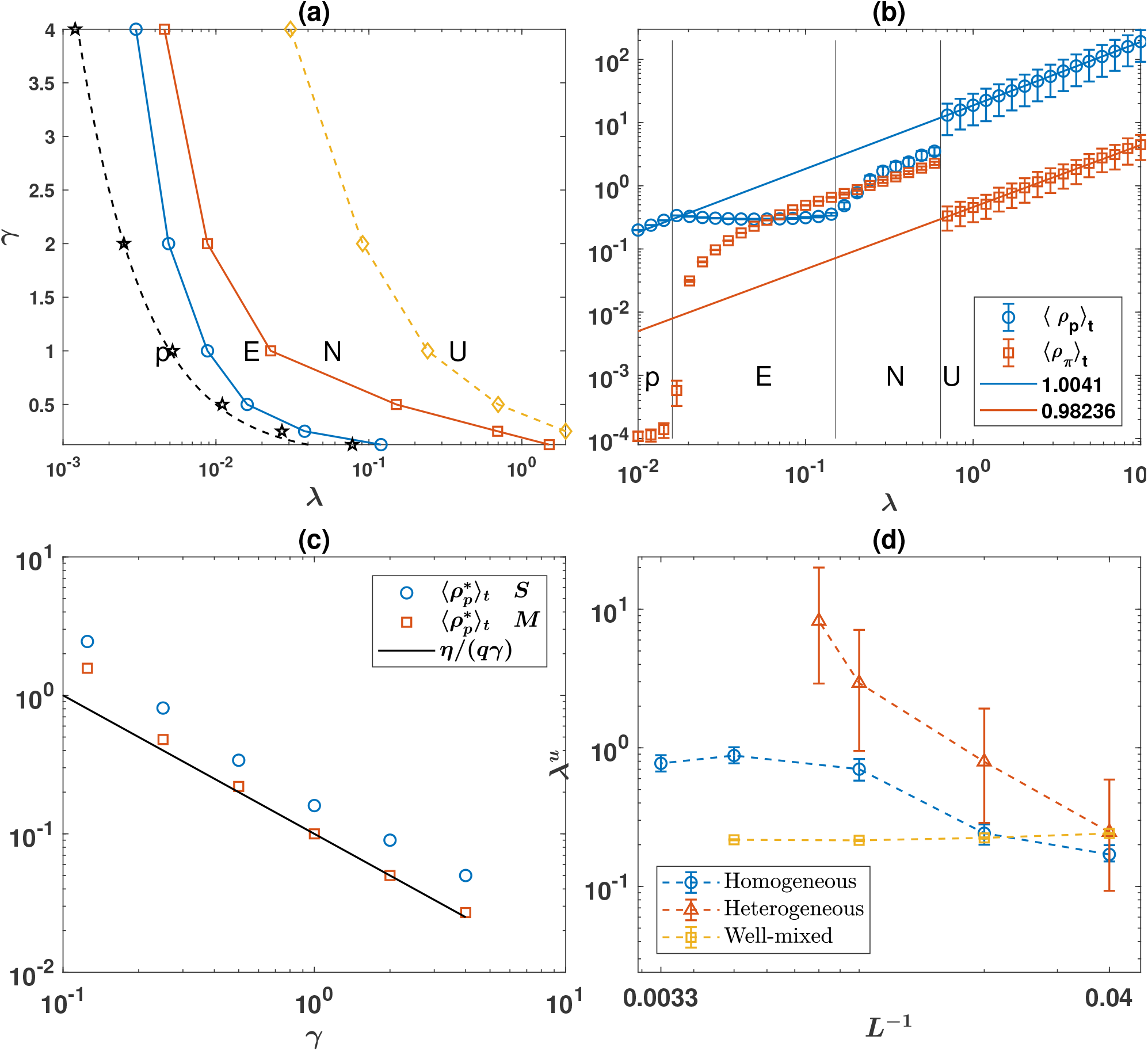
Phase diagram and phase transitions. The phase diagram of the model in the *λ − γ* plane (a), and the densities of prey and predators as a function of *λ* for *γ* = 0.5 (b). For too low resource regeneration rates, only preys survive (marked by the letter p). By increasing *λ*, the system shows a series of successive phase transitions to a phase where preys and predators coexist in equilibrium *E*, a non-equilibrium phase marked by traveling waves of preys and predators *N*, and a phase where the dynamics become unstable due to predators driving preys to extinction *U*. The dashed line, *λ*^*∗*^ = *η*^2^*/γq*, is a naive mean-field prediction for the phase transition above which predators survive, which is in good agreement with simulation results for a mixed population (black stars). (c): A\ naive mean-field argument by equating resource flow to predators *qρ*_*p*_*γ*, and resource consumption *η*, predicts the density of preys in the equilibrium phase to be equal to *η/*(*qγ*) (solid red line), in reasonable agreement with simulation results (blue circles for a structured population and red square for a mixed population). (d): The dynamics become unstable in a homogeneous environment for a high resource regeneration rate, manifesting the paradox of enrichment. The instability of the dynamics endures even for large system sizes (blue circles) in both structured (blue circles) and mixed (orange squares) populations. On the other hand, in a heterogeneous environment, the stability of the dynamic is restored for large system sizes (red triangles). In (a) and (b) the population network is a first nearest neighbor square lattice with von Neumann connectivity, fixed boundaries, and linear size *L* = 100. Parameter values: *η* = 0.05, *d* = 2, *γ* = 0.5, *q* = 0.5. Error bars in (b) are standard deviations divided by 2 to increase visibility.

By increasing *λ* beyond a second threshold (red squares), non-equilibrium fluctuations are set in a structured population. These fluctuations are derived when the density of predators becomes high enough such that the speed with which predators consume preys in a neighborhood is higher than the speed with which preys arrive in that neighborhood. As a result, the prey’s density field becomes inhomogeneous, and the dynamics of the model are derived by traveling waves of prey followed by fronts of predators (see the Supplementary Videos and the Supplementary Notes 1 and 2 for the dynamics of the model in this regime). A simple argument suggests this phenomenon occurs when predators become abundant enough to be able to deplete prey density field locally. That is when the number of preys consumed per site, which in the mean-field approximation is equal to *ρ*_*π*_*q*, becomes equal to the density of preys, *ρ*_*π*_*q* = *ρ*_*p*_ (see the Supplementary Note. 3).

By increasing *λ* further, above a third threshold (orange diamond), the dynamics become unstable. The instability of the dynamics occurs in both mixed and structured populations and results from the fact that in rich environments, prey can become too abundant to elicit a too fast exponential growth of predators, which acts faster than the regulatory feedback from prey density to predator density. As a result, the bottom-up regulation of the predator-prey system breaks, and the high predator density derives prey to extinction before preys’ low availability can regulate the density of predators. Investigation of the dependence of the value of *λ* above which the dynamics become unstable to system size in Fig. 1(d), shows that while for too small sizes, the instability threshold shifts to larger values in a structured population (blue circles), the system remains unstable even for very large system sizes. Investigation of other parameter values reveals the instability of the predator-prey dynamics in resource-rich environments is a characteristic feature of the model.

The instability of the predator-prey dynamic in resource-rich environments, coined the paradox of enrichment, is argued to result in simple models of predator-prey dynamics [24]. This theoretical prediction has also found some empirical support [17, 18, 23]. However, it is argued natural predator-prey systems can escape the paradox of enrichment using different mechanisms [23]. For instance, using theoretical models of predator-prey interactions, it is argued that spatial structure can stabilize predator-prey dynamics [25]. The model introduced here suggests that while spatial structure can improve the stability of the system by shifting the instability to richer environments (compare blue circles and orange squares in Fig. 1(d) for large system sizes), the instability of the dynamics endures on a spatial structure. However, as can be seen in Fig. 1(d) (red squares), resource heterogeneity provides a simple mechanism to restore the stability of the dynamics due to the fact that in a heterogeneous environment, resource-poor regions can provide a niche for preys in which they can survive due to the low abundance of predators and from time to time immigrate to resource-rich environments (See Section Methods).

Interestingly, in the non-equilibrium regime, the system appears to show a scaling regime in a homogeneous environment. However, due to the small range of the stable non-equilibrium region, it is not possible to convincingly argue for the existence of such a scaling regime in this region. By stabilizing the predator-prey dynamics, resource heterogeneity allows inspection of scaling in the stable regime and reveals that several variables, including prey density, predator density, prey’s density fluctuation, predators’ density fluctuation, preys birth rate, predator’s birth rate, and prey production, show scaling with *λ* in large densities (see the Supplementary Note. 5 and Supplementary Figure 8). The existence of scaling with *λ* suggests these variables show scaling as a function of each other. To see this, consider two generic variables, *v* and *w*, with a scaling relation, 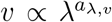 and 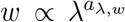. Then it is easy to see 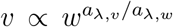. In Figs. 2(a) and 2(b), I plot the density of predators and prey production as a function of the density of prey for three different system sizes. As the system size increases, the dynamics stabilize up to larger values of *λ*. For *L* = 25, Both predator density and prey production show sublinear scaling with respect to prey density. For the unstable dynamics, the scaling is closer to linear (with a slope of *≈* 0.86), and in the stable regime, the scaling is close to a three-quarter scaling observed in predator-prey systems [11]. Another consequence of scaling relations of the model is Taylor’s power law. This can be seen in Fig. 2(c) and 2(d), where the density fluctuations of prey and predators as a function of their densities are plotted. Both preys and predators show a Taylor’s power law with an exponent of 0.5 in the equilibrium regime, indicating weak density dependence, and an exponent of 1 in the non-equilibrium regime, indicating highly aggregated dynamics and strong density dependence. As shown in the Supplementary Information, a sublinear scaling holds consistently for other parameter values and resource distribution (See the Supplementary Notes. 5 and 6)

**Figure 2:**
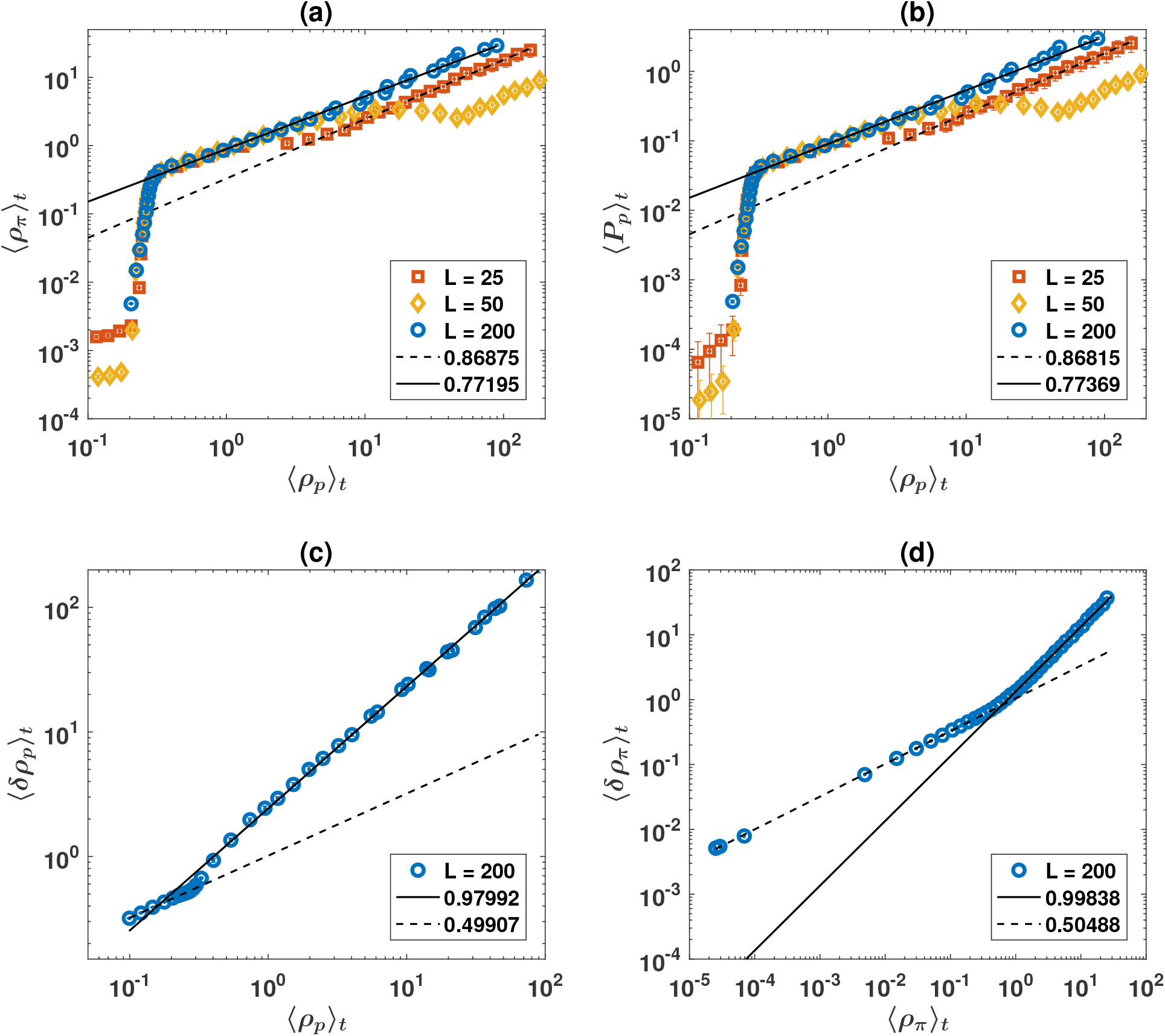
Predator-prey power laws. Predator density as a function of prey density (a) and prey production as a function of prey density (b) for three different system sizes and a linear distribution of resources are plotted. The system is unstable for large densities for small systems sizes (*L* = 25 and *L* = 50). The system shows a sublinear predator-prey and prey production-prey density scaling in large densities in both stable and unstable regimes. Both preys and predators show Taylor’s power law with an exponent close to 0.5 for small densities, indicating weak density dependence, and 1 for large densities, indicating a highly aggregated dynamic. Parameter values: *η* = 0.05, *d* = 2, *γ* = 0.5, *q* = 0.5. Errorbars are standard deviation divided by 5 to increase visibility.

A curious question is what mechanism drives the sublinear scaling? It is possible to derive an analytical form for prey production. Assuming there are *s*_*p*_ preys and *s*_*π*_ predators on a site, the expected number of preys caught by predators can be written as:

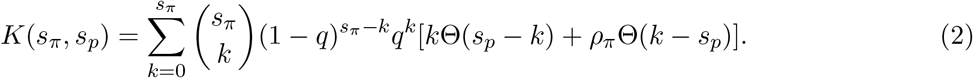

Where the theta function, Θ(*x*) is one if *x >* 0 and zero otherwise. To write down this expression, note that 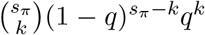 is the probability that *k* preys are caught. When *k* is smaller than the number of prey on the site, predators catch *k* prey. When k is larger than the number of preys, all the preys on the site are caught. Summing over *k*, we have the average number of preys caught on the site. Prey production is the average of *K*(*s*_*π*_, *s*_*p*_) over all the sites:

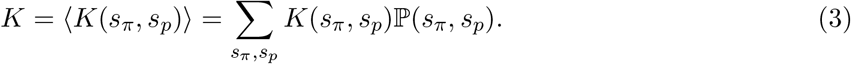

Where, ℙ(*s*_*π*_, *s*_*p*_) is the probability of observing *s*_*p*_ preys and *s*_*π*_ predators on a site. It is possible to approximate this expression by a simple production function:

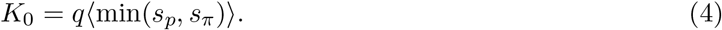

This expression is exact when the number of prey on a site is larger than the number of predators, and an approximation is involved when the number of prey is smaller than the number of predators. As can be seen in Fig. 3(a), *K* (solid blue) exactly describes prey production, and *K*_0_ (dashed red) shows a good agreement with simulation results. As we will shortly see, both these quantities show a scaling with the same exponent as prey production. Underlying these functional forms of prey production is a key assumption of the model, which I call the shelter effect: Predators get only one chance to attempt predation. Realistically, this feature can result from preys’ behavioral responses to an unsuccessful predation attempt as following an unsuccessful predation attempt, the prey can better avoid predation, for instance, by scattering around [30, 31], showing a higher vigilance [30–32], alarming the peers [30, 33, 34], or due to predator’s fatigue after an unsuccessful attempt. Importantly, this functional form is in stark contrast with the mass-action law functional response typically considered in Lotka-Volterra models [26–29], which can be written as the correlation function of prey and predators densities, ⟨*s*_*p*_*s*_*π*_⟩, times the probability of successful predation, *q*. As can be seen from Fig. 3(b), where mass action law functional response *q*⟨*s*_*p*_*s*_*π*_⟩ together with *K* and *K*_0_ are plotted, a mass action law functional response is a valid approximation is low densities, and as argued before, predicts an invariant prey density with respect to resource regeneration rate in low densities (*ρ*_*p*_ = *η/γq*). For larger densities, the shelter effect starts to play a role, and *K* and *K*_0_ can not be approximated by a mass action law. While a mass action law functional form is invariant to density fluctuations, the minimization procedure involved in *K*_0_, or the shelter effect reflected in *K* makes prey production sensitive to density fluctuations due to the fact that lower resource flow in places where preys are found in a small density (a density smaller than prey average density and predator density) is not compensated by a higher resource flow in prey rich places (where prey density is higher than both average prey density and predator density). As density fluctuation increases with density due to Taylor’s law, prey production decreases with density. This leads to a sublinear scaling of prey production and predator density as a function of prey density and, as we will see below, a prey-favoring life histories and bottom-up control of the food web in richer environments.

**Figure 3:**
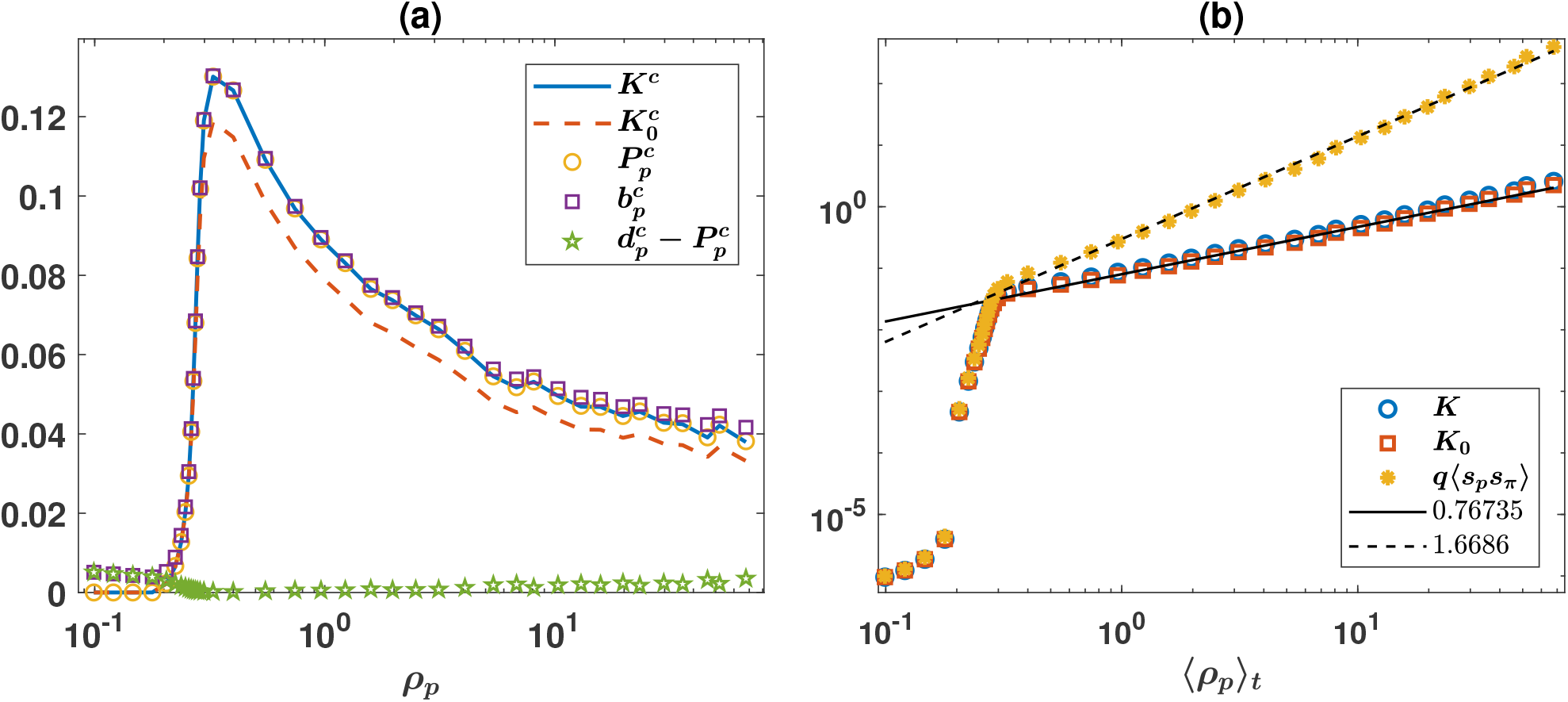
The shelter effect leads to a sublinear scaling. (a): 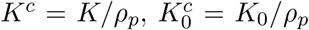, per capita prey production 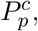, prey birth rate 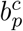, and natural death rate, 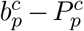, defined as death rate of preys due to going out of resource, as a function of prey density. *K* exactly describes prey production. (b) While *K* and *K*_0_ show scaling with prey density with the same exponent as prey production, the correlation function between prey and predator densities times *q, q(s*_*p*_*s*_*π*_), which determines prey production if mass action law holds, deviates from prey production in the non-equilibrium regime. The population resides on a first nearest neighbor lattice with von Neumann connectivity and fixed boundaries, with linear size *L* = 200. Parameter values: *η* = 0.05, *d* = 2, *γ* = 0.5, *q* = 0.5.

The model also sheds new light on the density dependence of life history strategies. In Fig. 4(a) and 4(b), I plot the distribution of lifespan of preys and predators for different values of *λ*. In the equilibrium regime, preys lifespan decays exponentially fast for short lifespans and shows a power-law decay for large lifespans. However, in the nonequilibrium regime, prey lifespan distribution develops a mode in large densities. This means that due to the shelter effect - the benefit of living in large groups - in large densities most of the preys are able to live for a characteristic lifetime before getting captured by predators. The lifespan distribution of predators on the other hand shows regular peaks. The *m*th peak occurs at *τ*_0_ + (*m −* 1)*d/η*, and corresponds to the predators who capture *m −* 1 preys during their lifespan. In the regime of large densities, the peaks for large *m* diminish. This means that in the non-equilibrium regime, predation becomes increasingly difficult due to the shelter effect and predators starve and die faster.

**Figure 4:**
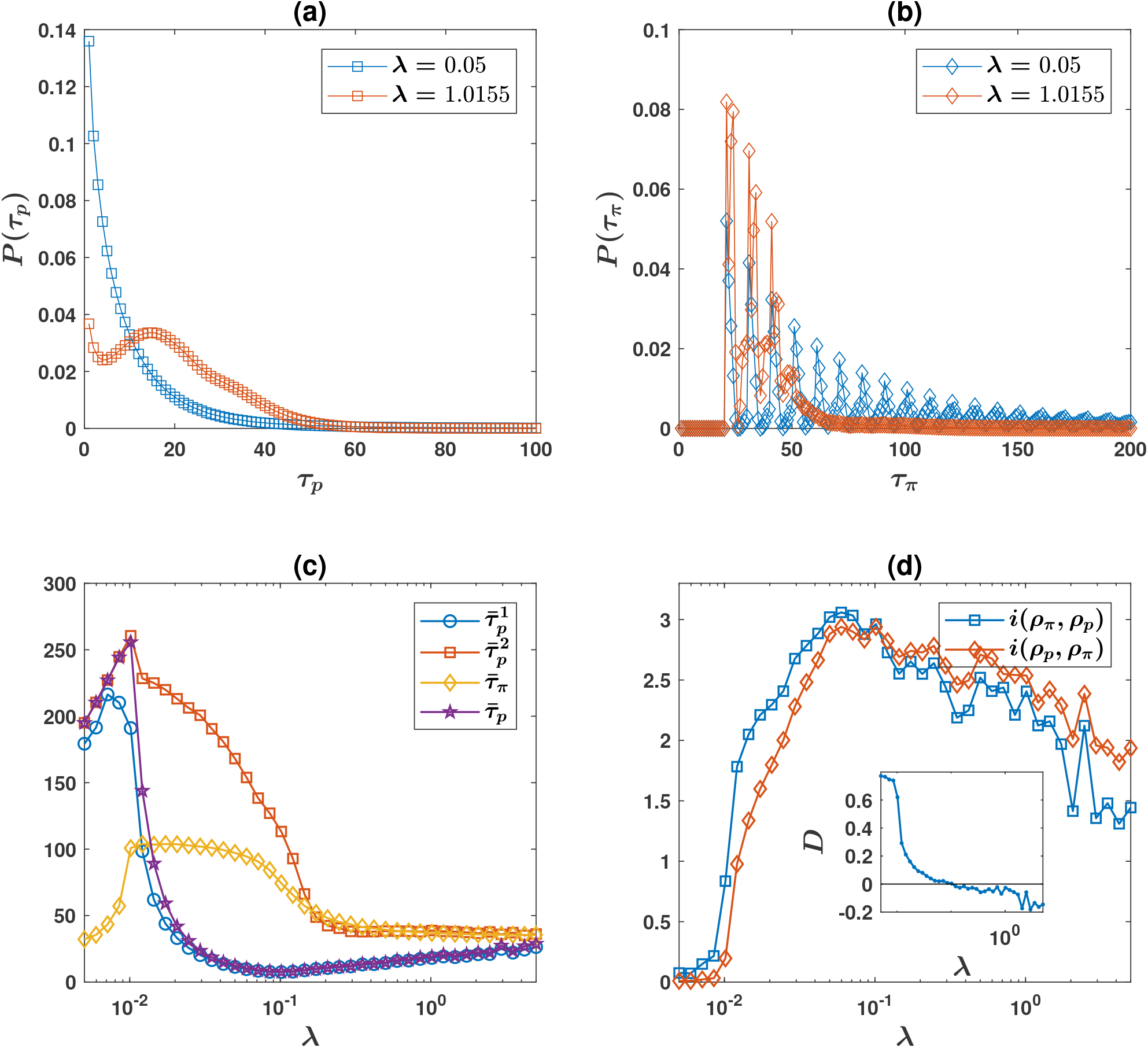
Life histories and food-web control. (a) and (b): The lifespan distribution of preys (a) and predators (b), in the equilibrium regime *λ* = 0.05 and non-equilibrium regime *λ* = 3.. While in small densities (equilibrium regime), most of the prey are captured rapidly and live short, in large densities (non-equilibrium regime), the lifespan distribution of prey develops a peak at a characteristic lifetime due to the benefit of living in large groups in large densities. Predators lifespan distribution shows peaks at intervals *d*_0_ + (*m −* 1)*d/η*. Each peak corresponds to predators who capture *m −* 1 preys during their lifetime. (c): Mean lifespan of preys 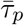, those preys captured by predators, 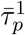, preys dying naturally 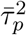, and the mean lifespan of predator 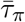. In small densities, resource availability does not affect predators’ lifespan significantly and decreases the mean lifespan of prey. In large densities, due to the advantage of living in larger groups for preys, the mean lifespan of preys increases, and predators’ mean lifespan slightly decreases by increasing density. (d) information flow from predator density to prey density, *i*(*ρ*_*π*_, *ρ*_*p*_) and from prey density to predator density *i*(*ρ*_*p*_, *ρ*_*π*_). Inset shows the causality direction of predators to preys defined by *D* = [*i*(*ρ*_*π*_, *ρ*_*p*_) *− i*(*ρ*_*p*_, *ρ*_*π*_)]*/*[*i*(*ρ*_*π*_, *ρ*_*p*_) + *i*(*ρ*_*p*_, *ρ*_*π*_)]. In small densities, predators drive the evolution of prey, suggesting a top-down control, and in large densities, preys drive the evolution of the system, suggesting a bottom-up control. The population network is a first nearest neighbor square lattice with von Neumann connectivity, fixed boundaries, and linear size *L* = 200. Parameter values: *η* = 0. 05, *d* = 2, *γ* = 0.5, *q* = 0.5.

Resource availability has contrasting effect on preys and predators’ mean lifespan as well (Fig. 2(c)). In small densities (equilibrium phase), predators’ mean lifespan do not show a significant dependence on density and their per-capita birth increases with increasing density. Preys’ mean lifespan decreases, and their per capita birth increases with increasing density. This shows that by increasing the resources, preys reproduce more and live shorter due to higher predator density and higher predation. The system shows different behavior in large densities (non-equilibrium phase): Preys mean lifespan increases with density while predators’ lifespan slightly decreases with density. The per capita birth rate of preys decreases with *λ* with an exponent of 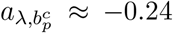 (see the Supplementary Note. 4 and Supplementary Figure. 7) and their lifespan increases. Predators’ per-capita birth rate increases with an exponent of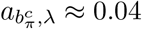, and their mean lifespan slightly decreases with *λ*. These observations imply that in rich environments, preys reproduce less and live longer, while predators reproduce more and live shorter. The difference in the lifespan of prey and predators, in turn, leads to the sublinear scaling of predator density as a function of prey density: As prey and predator per area birth rates scale almost linearly as a function of each other (see the Supplementary Note. 5 and Supplementary Figure. 8), longer lifespan of preys by increasing density leads to higher prey density in the system.

The analysis shows that predator-prey systems exhibit a top-heavy food web in small densities, and increasing resources favor predators in terms of both density and life histories. On the other hand, the food web is bottom-heavy in large densities. That is, higher resources are more advantageous for prey. This observation suggests that food-webs show different causality and control directions in small and large densities: In small densities, where equilibrium holds, food-webs exhibit top-down control and are controlled by predators, and in large densities, they exhibit bottom-up control and are derived mainly by prey. An examination of direction of causality based on flow of information from the time evolution of predators’ density to preys’ density, *i*(*ρ*_*π*_, *ρ*_*p*_) and information flow [35] from preys’ density to predators’ density *i*(*ρ*_*p*_, *ρ*_*π*_) shows this is indeed the case (4(b)). This finding suggests that the nature of the control in the food web results from prey and predators behavioral responses to each other. In small densities, preys get caught rapidly and higher resources leads to higher predation, higher predator density, and shorter lifespan of preys. Predators are the winners of competition in this regime and control the food web. In large densities, the shelter effect hands the control of the food web to preys, and the benefit of living in large groups give rise to the emergence of a characteristic lifespan of preys, which is further strengthened by increasing the density.

An interesting question is the origin of scaling. As I argue below, the answer to this question relies on the scale invariance of the group size distribution. To see this, in Fig. 5(a) and 5(b), I plot the distribution of group size, *s*, defined as the number of individuals in each lattice site, for different resource regeneration rates, and for, respectively, preys, *s*_*p*_, and predators, *s*_*π*_. The group size distribution of both prey and predator shows scale invariance with a lower cut-off for small group sizes, 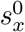, and an upper cut-off for large group sizes, 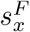 (here *x* stands for prey, *p*, or predator *π*). Postulating a simple scaling form *f* (*s*_*x*_, *λ*) *∼ λ*^*−β*^*f*_0_(*s*_*x*_*λ*^*−α*^), all the distributions for different values of *λ* collapse into a universal power-law *f*_0_. Besides, the upper cut-off, which is the largest group size, obeys a simple power-law relation with respect to *λ*, with an exponent, *ν*_*x*_ (Fig. 5(c) and 5(d)).

**Figure 5:**
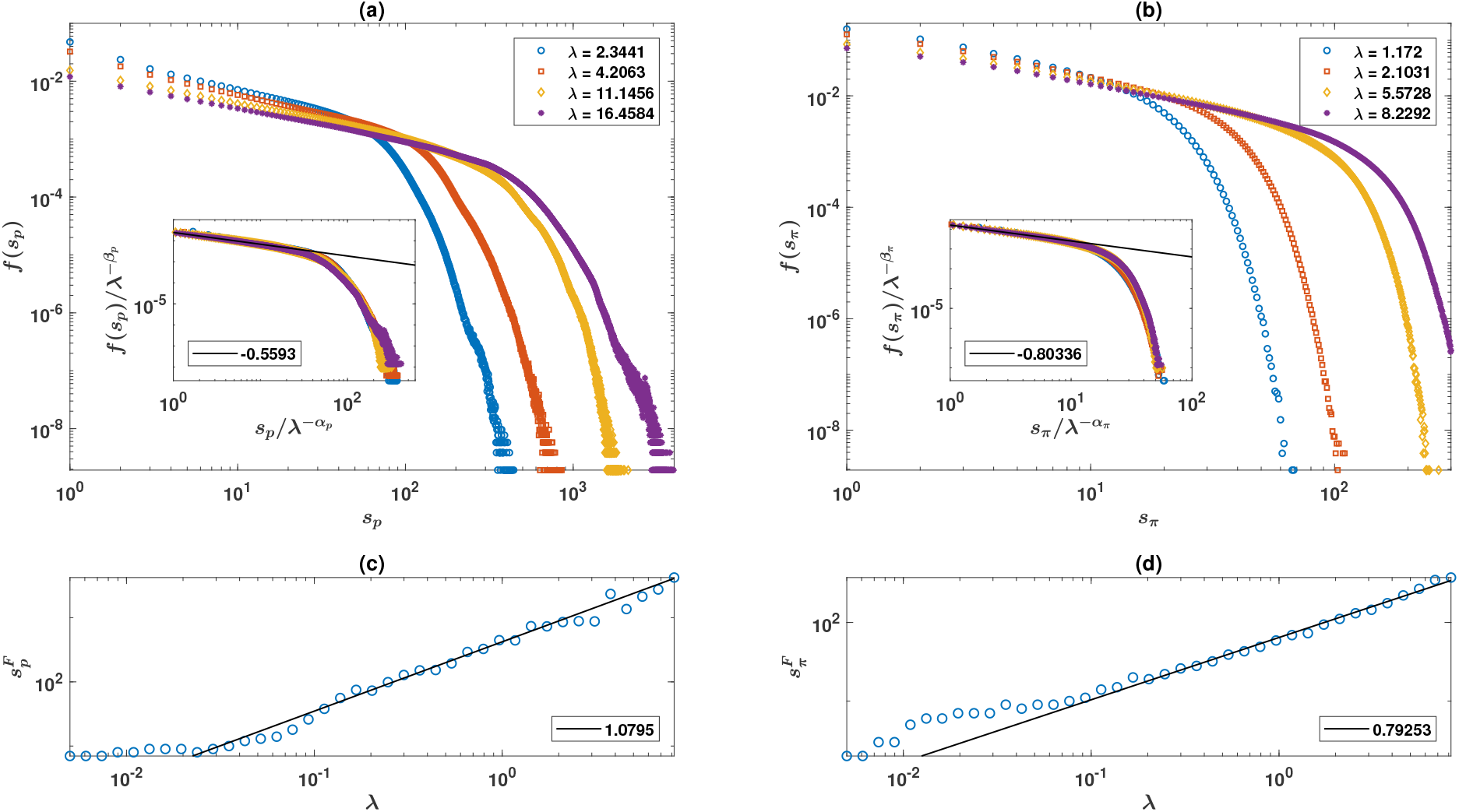
Scaling of the group size distribution. (a) and (b): Group size distribution of prey (a) and predators (b) show a power-law with a cut-off at large group sizes. Insets show data collapse for the re-scaled group size distributions. (c) and (d): Maximum group size shows scaling with *λ* at large densities. Groups size is defined as the number of individuals per lattice site. The population resides on a first nearest neighbor lattice with linear size *L* = 200. Parameter values: *η* = 0.05, *d* = 2, *γ* = 0.5, *q* = 0.5.

In the following I show these two conditions, that is the scale invariance of the group size distribution together with a scaling of the cut-off with resource regeneration rate lead to scaling of the species populations. To see how this is the case, consider the average density of species *x* in a given 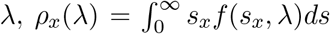 (For discrete group size, *s*, we have 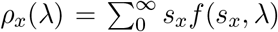. One can use Euler-Maclaurin theorem to approximate this sum by an integral). Assuming the lower cut-off is zero, and using the simple scaling form we have 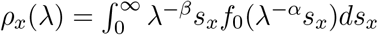. Using the fact that *f*_0_ shows a sharp cut-off, the infinity in the upper bound of integral can be replaced by the cut-off, 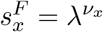 and assuming *f*_0_(*s*_*x*_) has a power law form, 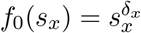, we have:

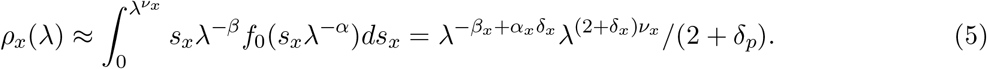

Thus, 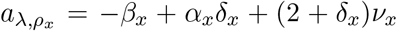. Using similar calculations, it is possible to shows that the exponent of the *n*th moment of population density of species 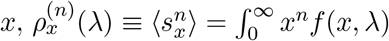, is equal to 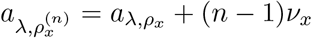. Finally, noting that in the stationary state the prey production should balance predator consumption, *K*(*λ*) = *ηρ*_*π*_(*λ*), we get the same exponent for prey production, 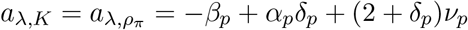.

Using the normalization of the probability density function, it is possible to derive approximate equalities for the exponents. Neglecting the mass of the probablity density function above the upper cut-off and taking the lower cut-off equal to zero, we have 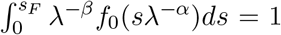, from which one can derive *−β*_*x*_ + *α*_*x*_*δ*_*x*_ = *−*(1 + *δ*_*x*_)*ν*_*x*_. This implies 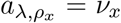, which predicts a Taylor’s power-law with an exponent of one, 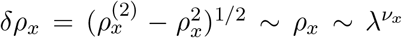, that is a highly aggregated dynamics observed in the non-equilibrium regime. The best estimates of 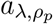 and 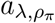 based on a log-log fit are respectively *≈* 1.19 and *≈* 0.91 (see the Supplementary Figure 8), and the best estimates for *ν*_*p*_ and *ν*_*π*_ are, respectively, 1.08 and 0.79 (Fig. 5). A linear fit in a log-log plot suggests *δ*_*p*_ = 0.57 and *δ*_*π*_ = 0.80 (Fig. 5). Upon inspection, *α*_*p*_ = 1.1 and *β*_*p*_ = 0.95 leads to a reasonable data collapse, which gives 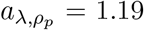. For predators, *α*_*π*_ = 0.9 and *β*_*π*_ = 0.95, which gives 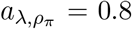. The exponents continue to satisfy these equalities with an error of up to about 10 percent for other parameter values, which can be attributed to taking the lower cut-off equal to zero, and neglecting the mass of integral above a upper cut-off, as well as errors involved in a determination of exponents using a log-log plot. As we have seen here, a simple model of predator-prey interactions gives rise to a scale-invariant ecosystem, with a scale-invariant distribution of group sizes. This, in turn, can give rise to a rich set of scaling laws. Notably, the theory developed here is rather general. Its only assumptions are that the distribution of group sizes obeys a power-law relation with a cut-off proportional to a power of resource availability. These assumptions appear to hold in many species and ecosystems. For instance, it is argued that group size distributions are truncated power laws [12, 13, 15, 16], or power-laws with exponential decay, and the universality of these distributions are argued [14]. In addition, an increasing relation between density and group size is often observed in animal groups [36–38]. Our result suggests a universal power-law group size distribution can simply result from predator-prey interactions with a universality class that is different for prey and predators. Besides, maximum group size shows a sublinear power-law relation with respect to density. As we have shown, these observations can lead to a rich set of scaling laws, as long as different species coexist and interact in an ecosystem. Importantly, this argument is independent of the nature of the interactions and extends well beyond predator-prey interactions.

While the existence of different phases in predator-prey systems has been reported experimentally [23], a theoretical framework showing how these phases can appear and relate to ecosystem functions and structures was lacking. The model introduced here can provide such a framework and shows that a simple model of predator-prey interaction predicts different phases to be at work in such systems. This finding suggests inconsistencies between past empirical results regarding predator-prey dynamics can simply result from looking at this system in different physical regimes. The existence of different phases in food webs also has interesting implications for one of the most fundamental questions in ecology: the question of whether food-webs exhibit top-down or bottom-up control? [20–22]. The model shows a top-down control results in small densities where the system shows equilibrium and non-aggregated dynamics, and bottom-up control results in large densities with the onset of collective motion of prey and predators with highly aggregated dynamics, where the shelter effect starts to play a role, and a characteristic lifespan of preys emerges. This finding shows preys’ collective responses to predators have important consequences for fundamental ecological patterns and determine the nature of control in food webs. This observation shows the importance of going beyond simple Lotka-Volterra models by taking the behavioral responses of prey and predators into account [39] and how these behavioral factors can underlie fundamental ecological patterns [40].

## Methods

### The simulations

At each time step of the simulation, at first, preys consume resources. For this purpose, the resources available on each site are equally divided among the prey visiting that site. Then, predators are given a chance to capture prey. For this purpose, each predator on the site is given a chance to attempt to capture a prey. The predator is successful with probability *q*. This is continued until either all the prey on the site are captured or all the predators on the site attempt capturing a prey. In the next stage, preys and predators reproduce and move. For this purpose, those with an internal resource above *d* reproduce an offspring, and the resources are divided equally among the agent and its offspring. After reproduction, agents move to one of the neighboring sites chosen at random. Agents pay for the metabolic cost, *η*, and those with internal resources below 0 die out. Finally, the resources available on each site ***r*** increases by a rate *λ*(***r***). If it happens that the population of one of the species goes to zero, assuming there is migration, one individual of that species with internal resource 1 is added to a random location on the lattice. At the beginning of the simulations, the individuals are distributed in random locations on the lattice. After checking that initial conditions does not affect the equilibrium dynamics, all the simulations start with 100 prey, 10 predators, and initial internal resource equal to 1.

### Resource distribution

In the case of homogeneous resource distribution, the primary resource regeneration rate is taken to be a site-independent constant *λ*(***r***) = *λ*. For a linear resource regeneration rate, the primary resource distribution is given by *λ*(*r*) = 2*λx/L*, where *x* is the *x*-component of the position, ***r, r*** = (*x, y*). The factor of 2 ensures the average resources regeneration rate in the system to be equal to *λ*, and thus comparable with the homogeneous case.

In the Supplementary Information, I have also considered a two-patch environment or binary resource distribution. In the binary resource distribution case, the environment is divided into two patches, a poor patch, extending from *x* = 0 to *x* = *L/*2, with a constant resource regeneration rate *λ*_0_, and a rich patch, extending from *x* = *L/*2 to *x* = *L* with a resource regeneration rate *λ*.

### Variable definition

Prey density, *ρ*_*p*_ (predator density, *ρ*_*π*_), is defined as the number density of preys (predators) defined as the total number of preys (predators) in the population divided by the area. In order to define prey and predator density fluctuation (*δρ*_*p*_ and *δρ*_*π*_) we first define the density field as the number of preys living on a lattice site (*x, y*), *s*_*p*_(*x, y*). The density fluctuation of preys is the standard deviation of this quantity 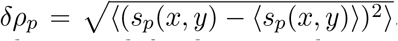, calculated over all the lattice sites. The density fluctuation of predators is defined in a similar way.

Prey and predator per area birth rate (*b*_*p*_ and *b*_*π*_) are defined as the average number of births per lattice site, derived by dividing the total number of births in a time step by lattice size. Per capita birth rates (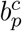 and (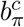) are defined as the average birth rate of a prey (predator) per time step. These are derived by dividing the per area birth rate by density. Prey and predator per area death rates (*d*_*p*_ and *d*_*π*_) are defined as the average number of deaths per lattice site. To calculate these, we divide the total number of deaths in a time step by lattice size. Per capita death rates (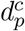 and 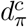) are defined as the average death rate of a prey (predator) per time step. These are derived by dividing the per area birth rate by density. Prey per area production (*P*_*p*_) is the number of preys captured by predators per lattice site. Prey per capita production, 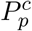 is defined as per area prey production divided by prey density.

Group size of preys, *s*_*p*_ and predator, *s*_*π*_ is defined as the number of individuals in a lattice site. Maximum groups size is defined as the maximum group size observed in the stationary state of the simulation ran for *T* time steps (*T* = 20000 in Fig. 5 and the fist 1000 time steps are discarded as non-stationary period). The lifespan of preys and predators is defined as the number of time steps that a prey or predator lives. Lifespan distributions are calculated based on a simulation run for *T* time steps and after the system reaches stationarity (*T* = 15000 in Fig. 4 and the first 1000 time steps are discarded as non-stationary period).

### Mean field results

To derive an approximate relation for the phase transition, note that assuming mass action law holds (which is a valid assumption in the regime of low densities), the average flow of resources into predators is equal to *γqρ*_*p*_, which should become larger than *η* at the transition point. Assuming that at the transition point, the prey’s density remains close to its stationary value in the absence of predators, *ρ*_*p*_ = *λ/η*, (based on the fact that predators’ density is close to zero at the transition), we arrive at the naive mean field prediction, *λ*^*∗*^ = *η*^2^*/γq*. This prediction is plotted by a dashed black line in Fig. 1(a), which provides a lower bound for the estimated transition line from simulations in a structured population (blue circles), and is in high agreement with simulation results in a mixed population (black stars). The deviation from the mean-field prediction in a structured population can be understood by noting that predators deplete the prey density in their neighborhood. Consequently, a larger prey density than predicted by the naive mean filed argument is necessary to sustain predators in the system.

To approximate the density of preys in the equilibrium regime, note that resource gain by a predator, *ρ*_*p*_*γq* and resource consumption, *η* should balance in the stationary state, which gives *ρ*_*p*_ = *η/*(*γq*). This argument grasps the overall dependence of *ρ*_*p*_ on *η, γ*, and *q*. This can be seen in Fig. 1(c), where the stationary density of preys at the onset of transition for different values of *γ* in a structured (blue circles) and mixed (red squares) population is compared with *η/*(*γq*) (black line). In a structured population, a deviation from simulation results is observed due to the fact that the local depletion of the prey density field as a result of predator presence which requires a higher prey density in this regime than that predicted by the naive mean-field argument, is not taken into account in the mean filed argument.

### Stabilization of the predator-prey dynamics by resource heterogeneity

To see how resource heterogeneity stabilize predator-prey interactions, consider a heterogeneous environment, where the resource regeneration rate on a site (*x, y*) on the lattice is given by *λ*(*x, y*) = *λ*2*x/L*. Thus, the average resources in such an environment are equal to *λ*. The value of *λ* above which the system becomes unstable for different inverse system sizes, *L*^*−*1^ is plotted in Fig. 1(d) (red squares). For too small system sizes, resource heterogeneity fails to stabilize the dynamics. This is due to the fact that for too small system sizes, there is effectively no separation between resource-poor and rich regions, and predators can immigrate from rich to poor regions and derive preys to extinction. However, as system size increases, fewer predators penetrate the poor environment. Consequently, the system becomes stable for larger and larger average resource regeneration rates, *λ*, in a heterogeneous environment. Heuristically, the system size required for the stability of the system is determined by the distance predators can penetrate into the stable region. Upon birth, a predator has a resource comparable to *d* and it survives for approximately *d/η* time steps without catching prey. With a linear resource distribution, the dynamic becomes unstable at a position *x*^*u*^ = *Lλ*^*u*^*/*2. From this position, predators can penetrate into the poor environment up to a depth approximately equal to *x*_0_ = *Lλ*^*u*^*/*2 *− d/η*. The dynamic is stable if *x*_0_ is significantly larger than zero. Equating *x*_0_ to zero, we get a characteristic system size, *L*^*u*^ = 2*d/*(*ηλ*^*u*^), which for the base parameter values is approximately equal to 60. In agreement with simulation results, this heuristic argument suggests that the dynamics are stable when the system size is a few times larger than 60. Although I have cast the heuristic argument in terms of linear resource distribution, The argument grasps the mechanism according to which resource heterogeneity can stabilize the predator-prey dynamics for sufficiently large system sizes, irrespective of the details of resource distribution.

### Direction of control

To examine the direction of control in Fig. 4(d), I have used direction of information flow between the time series of the density of preys and predators [35]. Information flow from the density of preys to predators is defined as 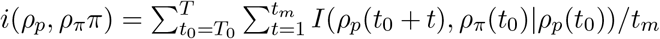. Here, *I* stands for conditional mutual information, and *ρ*_*p*_(*t*) (*ρ*_*π*_(*t*)) is the density of preys (predators) at time *t*. In Fig. 4(d), *t*_*m*_ is taken equal to 4, and a simulation ran for *T* = 15000 time steps is used after discarding the first *T*_0_ = 1000 time steps. The results are robust with respect to the choice of these parameters. The direction of information flow can be defined as the normalized difference of the information flow from the density of predators to preys and vice versa: *D* = [*i*(*ρ*_*π*_, *ρ*_*p*_) *− i*(*ρ*_*p*_, *ρ*_*π*_)]*/*[*i*(*ρ*_*π*_, *ρ*_*p*_) + *i*(*ρ*_*p*_, *ρ*_*π*_)] [35].

## Acknowledgment

The author is thankful to Ian Hatton, Onofrio Mazzarisi, Matteo Smerlak, Anton Zadorin, and other members of the structure of evolution group for insightful discussions. The author acknowledges funding form Alexander von Homboldt Foundation in the form of a Sofja Kawalevskaja award and the Deutsche Forschungsgemeinschaft (German Research Foundation) under Germany’s Excellence Strategy-EXC 2117-422037984 during parts of this research.

